# TTSBBC: Triplex Target Site Biomarkers and Barcodes in Cancer

**DOI:** 10.1101/2024.01.26.577471

**Authors:** Maya Ylagan, Qi Xu, Jeanne Kowalski

## Abstract

The technology of Triplex-forming oligonucleotides (TFOs) provides an approach to manipulate genes at the DNA level. TFOs bind to specific sites on genomic DNA, creating a unique intermolecular triple-helix DNA structure through Hoogsteen hydrogen bonding. This targeting by TFOs is site-specific and the locations TFOs bind are referred to as TFO target sites (TTS). Triplexes have been observed to selectively influence gene expression, homologous recombination, mutations, protein binding, and DNA damage. These sites typically feature a poly-purine sequence in duplex DNA, and the characteristics of these TTS sequences greatly influence the formation of the triplex. We introduce TTSBBC, a novel analysis and visualization platform designed to explore features of TTS sequences to enable users to design and validate TTSs. The web server can be freely accessed at https://kowalski-labapps.dellmed.utexas.edu/TTSBC/.

## INTRODUCTION

Triplex-forming oligonucleotides (TFOs) bind to the major groove of genomic DNA, termed TFO Target Sites (TTS), forming intermolecular triplexes via Hoogsteen hydrogen bonding (1,2). TTSs are poly-purine duplex DNA with binding influenced by longer lengths, higher G-content, and fewer pyrimidine interruptions (3-7).

Triplex formation is a promising cancer treatment as an anti-gene technique, capable of specific DNA sequence targeting (2,8,9). TFOs can direct sequence-specific DNA damage whether they are bound to another compound or by themself (2,10), and they can facilitate site-specific DNA repair synthesis and mutagenesis, likely via the nucleotide excision repair (NER) mechanism (2,11). TFOs also modulate transcription and inhibit replication (9). Common uses of TFO technology includes gene modulation by blocking transcription and gene modification through mutagenesis and homologous recombination (2,9,12). TFO binding is site-specific (13), leading to differential responses in cells with varying TTS abundance, which can be exploited to target cancer cells characterized by genomic instability (14). The site-specific nature of TFO binding to TTSs plays an important role in predicting the triplex formations in the genome.

Though there have been several databases of TTS (15,16), their exploration for potential as markers in cancer treatment strategies has been limited due to the lack of a corresponding analysis platform. We present TTSBBC as a webserver analysis tool for researchers investigating the role of TTSs in cancer and other genetic diseases within an accessible platform for analyzing and visualizing TTSs using their feature contexts.

## MATERIAL AND METHODS

### The web application architecture

The web application is developed using Shiny R. The front end of the application is extended with HTML, Cascading Style Sheets (CSS), and JavaScript. TTSBBC is deployed on Amazon Web Service (AWS). The deployment uses AWS Fargate with containerization. AWS Elastic Load Balancer (ELB) serves as a traffic controller to ensure visiting experience and scalability. TTSBBC accesses a secured MySQL database containing all possible TTS mappings and features that have been precalculated for improved user experiences. This website is free and open to all users and there is no login requirement.

### Data sources

The TTS sequences were downloaded from two sources, the Triplex-forming Oligonucleotide Target Sequence Search engine (TFO-Search) (15) and from TTSMI (16). These datasets denote human genome TTSs that comprise 1,297,671 TTSs from TFO-Search (17) and 36,276,455 TTSs from TTSMI (16), of which 384,089 overlapped between them. A database of 31 published, experimentally-validated TTSs was curated and is available to explore within the application which provides associated information and citations. Cancer cell line copy number segment data was sourced from DepMap (18,19) and filtered for amplifications, with names converted to cancer cell line encyclopedia nomenclature. The human genome reference sequence and annotation used was Gencode v44 (20).

### Data Pre-Processing

Each TTS sequence was aligned to Gencode v44 using Bowtie (16,21). When a TTS mapped to multiple genomic locations, all mappings that were exact matches were retained. For each TTS, sequence level features, length and associated base content, were also calculated. While each TTS is characterized by a feature set, their mappings to the genome can be one-to-many. The number and location of TTS mapping(s) were determined along with features as unique to a given TTS sequence. This precomputing enhances the analytical efficiency and user experience of TTSBBC.

A consensus gene element dataset was generated from the Gencode v44 GFF file. Promoters, introns, and exons were determined for all transcripts, then filtered to one representative transcript per gene. Introns were defined as unannotated regions (i.e. not exons, cds, or start/stop codons) while promoters were identified as 2Kb upstream of the start codon (15). In parallel, for a given gene, representative transcripts were selected based on this criteria in order: basic annotation, longest transcript (15), most exons, most annotated features, greatest support, and Ensembl canonical. In the case of ties, which occurred in less than one-percent of all genes, one transcript was randomly selected after manual review. To combine representative transcripts and gene elements, the gene elements were filtered for selected transcripts, then for exons, introns, and promoters to use as a dataset to annotate TTS mappings.

### Biomarker: TTSs as multi-gene targets

The accuracy and precision of a TTS relate to the user’s specified query region(s). For each TTS within these regions, both accuracy, indicating the degree of on-target mapping, and precision, reflecting the degree of overlap with query regions, are assessed. Collectively, these measures evaluate the TTS’s on and off-target mappings in the human genome and are defined as follows:

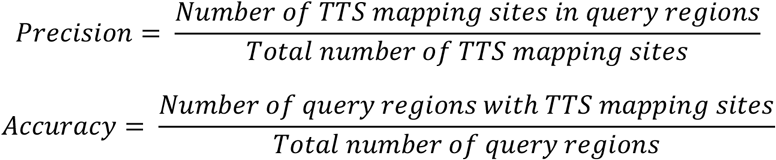

### Barcoding: Sequence-informed signatures

The link between TTS sequences, their genomic mappings, and sequence features are used to derive a barcode that may be further linked with a gene signature. For this task, each TTS’s genomic mappings were intersected with genomic regions to define a unique gene signature. Barcoding TTSs organizes them into gene signatures based on binding-relevant features such as length, G-content, and pyrimidine interruptions (3-7). Barcoding is key to performing a detailed analysis of TTS sets.

Barcoding of TTSs involves categorizing them based on key features, with each Barcode representing a combination of these traits. G-content and length are divided into two groups: above (G1/L1) or below (G2/L2) the median of that data source. Specificity is high (S+) for TTS sequences with multiple genomic mappings, and low (S1) for single mappings. Pyrimidine interruptions are classified as high (P+) if present, and low (P0) if absent. The presence of a G-quadruplex motif (22) is indicated as Q+, and its absence as Q0. For instance, a Barcode G1_L2_S1_P+_Q0 signifies a TTS with high G-content, shorter length, one mapping, at least one pyrimidine interruption, and no G-quadruplex motif. Using this barcoding system, all TTSs were organized according to the aforementioned traits.

Barcode signatures are representative of genes, TTSs, and TTS features; these components can be stratified (Supplemental Figure 1). This stratification illustrates the composition of Barcode signatures from various TTS gene signatures by data source. For instance, a desirable Barcode G1_L1_S+_P0_G0 contains TTSs with varying TTS gene signature sizes demonstrating sequence homology across the genome. Within each barcode, TTS gene signatures may be combined to derive barcode-level gene signatures (Supplemental Data 1). Barcoding adds a new dimension to genomic annotation, enhancing the interactive understanding of gene targeting using TFOs. This serves as a valuable reference for researching genes with specific TTS characteristics, aiding studies in TFO formation and advancing our knowledge of anti-gene therapy in cancer.

## RESULTS

### Overall design of TTSBBC

The TTSBBC web server features two main modules, as depicted in Figure 1. The first, “findTTS,” includes “Biomarker” and “Barcode” submodules and accepts various genomic region inputs. “Biomarker” calculates the accuracy and precision of a TTS, assessing on- and off-target mappings. “Barcode” evaluates the sequence-level features of TTSs for binding. The second module, “myTTS,” enables users to explore their TTS sequences, focusing on both features and genomic mappings.

**Figure 1.**
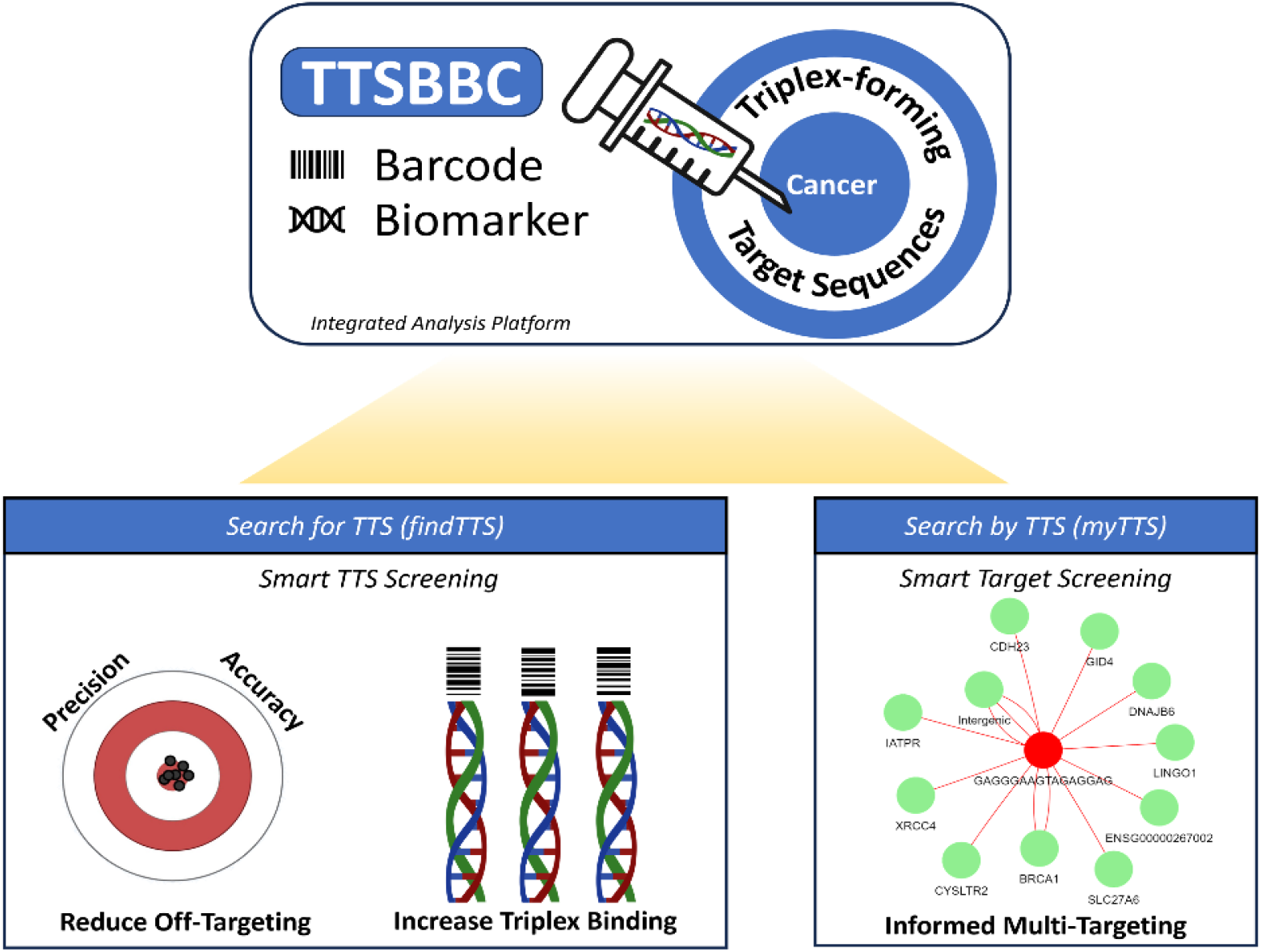
Overview of the TTSBBC analysis platform. TTSBBC is a platform for analysis of TTS sequences. The platform is designed with two paths of use in mind. Searching for TTS and searching by TTS. When searching for TTS, users are provided with analyses that guide the user to have smart screening of TTSs by reducing off-targeting and helping increase the triplex binding. When searching by TTS, users are provided with a smart screening of TTS targets and visualizations to help inform the multi-targeting of the a given TTS.

### Input format options

The web server offers various input methods. For “findTTS,” users can input genomic regions in several ways: using built-in cancer-related gene sets (23-30), selecting amplified segments from CCLE cell lines (31), manually entering genes, or uploading custom genomic regions via a csv file. The “myTTS” module accepts a TTS nucleotide sequence as its query.

#### findTTS module enables TTS discovery for a genomic region query

The “findTTS” module offers a comprehensive analysis tool for screening TTSs based on their accuracy and precision to target user-defined genomic regions (Figure 2). The “Biomarker” submodule enables researchers to compare TTSs’ accuracy and precision across different regions, identifying genes with varied on- and off-target mappings. It includes a network visualization to show which genes are targeted by TTSs within the input regions. The “Barcode” submodule categorizes TTSs into specific Barcodes based on sequence-level features. This module provides visual representations of TTS Barcode distributions, making analysis more intuitive and effective (Figure 3A). It also shows spatial relationships of TTSs, particularly those with the S+ Barcode, allowing detailed exploration of their genomic regional density (Figure 3B). Additionally, “findTTS” offers custom TTS Barcoding through k-means clustering based on numerical features, helping users identify optimal TTSs for their specific needs (Figure 3C). This feature supports deeper investigation into how sequence features influence TTS formation and function, enhancing user understanding of TTS characteristics and their impact.

**Figure 2.**
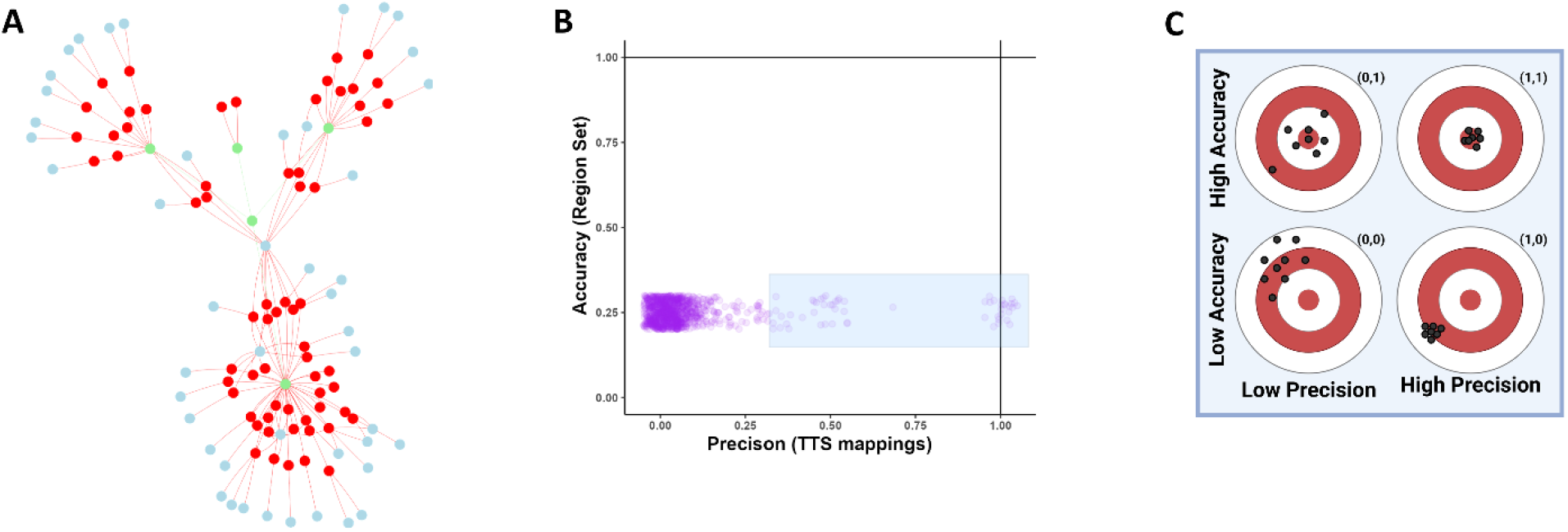
TTS Biomarker assessment using accuracy and precision. **A**, Network of TTS Mappings. The node colors in red are TTS sequences, in blue are off target genes, and in green are query regions. In the app users can zoom in and see the names of each node. The off-target mappings that are not within any gene region are connected to a blue node labeled intergenic. **B**, TTS accuracy and precision scatterplot. On the x-axis is precision, and y-axis is accuracy, both on a scale from 0-1. This scatterplot allows users to select TTSs as shown by the blue rectangle selection. **C**, A visual representation of accuracy and precision in relation to a target. Greater accuracy is close to the bulls-eye, how “on-target” a TTS is, and greater precision is a greater spread, how consistent and able to target all regions a TTS is. **D**, A table to aid the users in the interpretation of accuracy and precision values to relate figures B and C.

**Figure 3.**
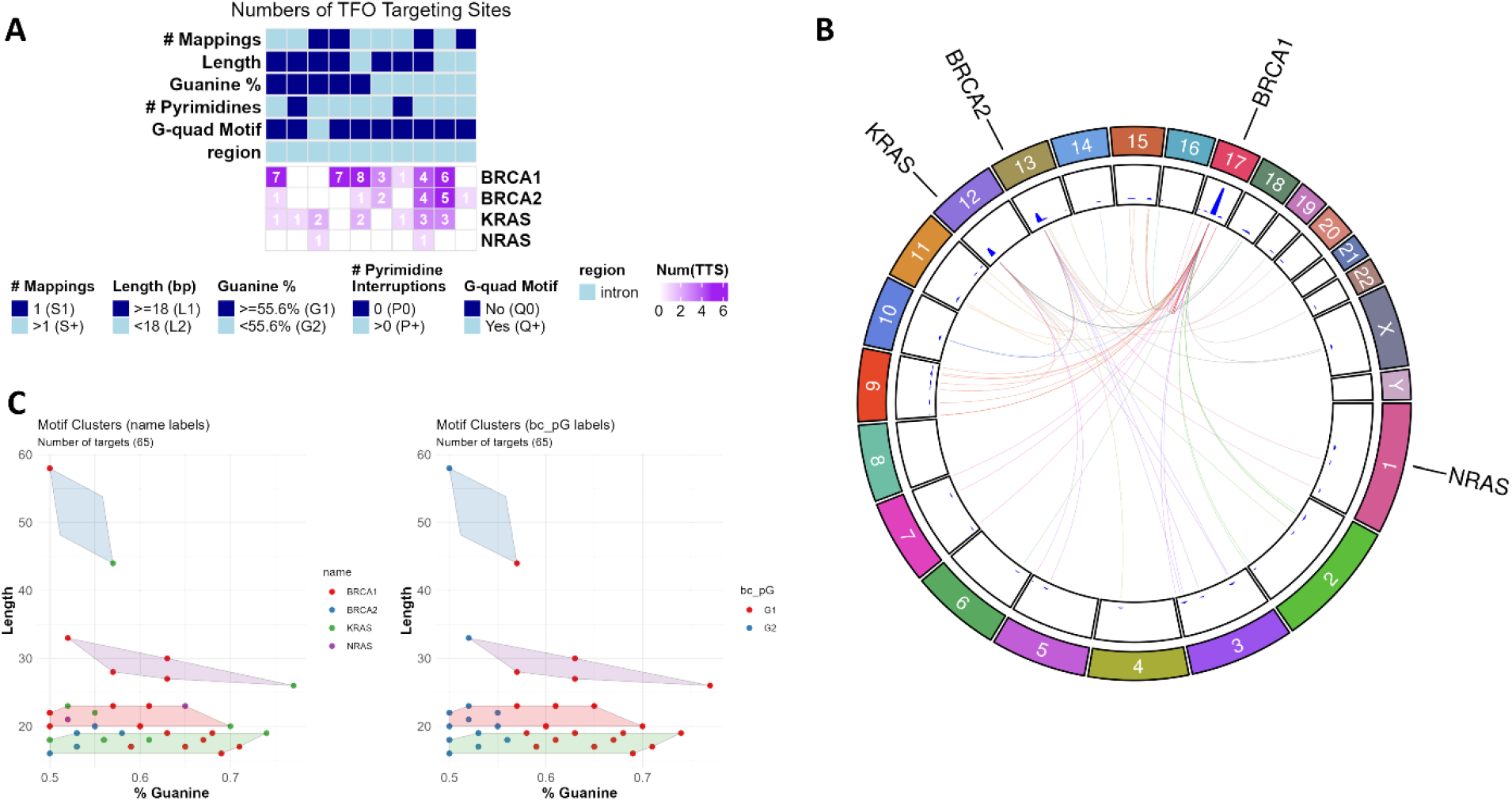
TTS barcode assessment of sequence level features. **A**, Heatmap of TTSs and their features in relation to their mapping to query regions. TTS barcodes are visualized as annotations above the heatmap to demonstrate the diversity of sequence features for TTSs that are within certain query regions. **B**, Circos plot denoting the locations the selected TTSs map to that are not within the query regions (labeled outside the chromosome ring). The TTS is linked to other regions it maps to, and the link is colored by the destination chromosome. **C**, Custom Barcoding. Clustering can be done on a pair of numeric features for users to create their own barcodes and find a set of TTSs that are most applicable to their work and use cases.

#### myTTS module enables TTS validation and investigation

The “myTTS” module focuses on TTS sequence analysis, enabling users to perform smart screening of input sequences. It identifies mappings for a given TTS and displays a network of genes connected to it. Additionally, the module offers detailed sequence-level features and characteristics, crucial for assessing TTS binding properties.

#### Case Study #1: Validation and Discovery of TTSs targeting HER2 in Breast Cancer

The recent study by Tiwari et al. (32) showed the enhanced ability of TTS’s to target Her2 as compared to Trastuzumab in breast cancer cell lines. Using their TTSs (with “myTTS” module), and cell lines (with “findTTS” module) we demonstrate the utility of our application. The “myTTS” module validated four TTS sequences (HER2-1, HER2-205, HER2-40118, HER2-5922) targeting the ERBB2 gene, with HER2-1, also noted in other literature (16), serving as a positive control (Figure 4). TTSBBC offers dual methods for TTS discovery: 1) targeting amplified cell line segments or 2) targeting genes. The cell line approach focuses on increased TTS copies for differential triplex formation and DNA damage in cells with amplified regions (32,33). Targeting genes directly is similarly effective, especially in cases of gene amplification. This dual approach showcases TTSBBC’s versatility in TTS analysis.

**Figure 4.**
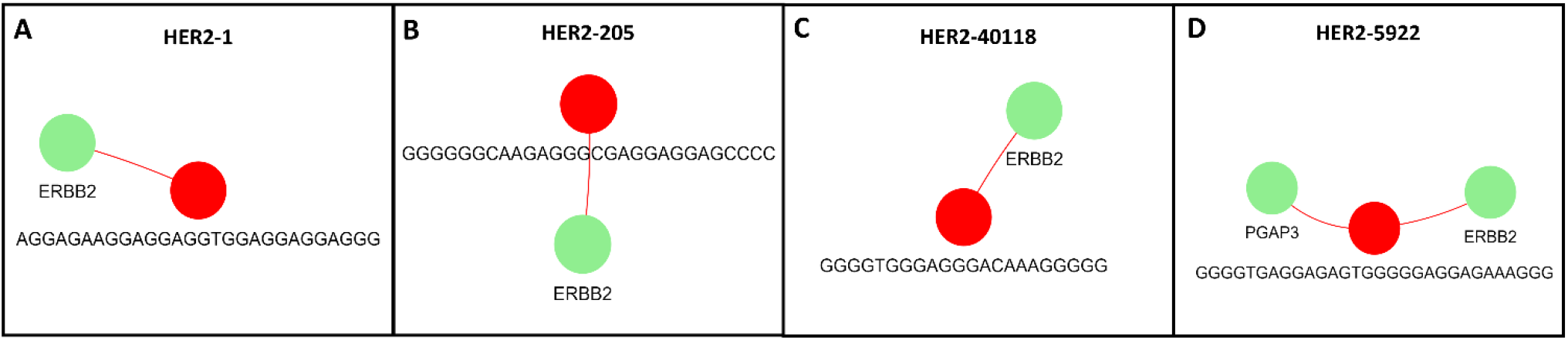
Network analysis of TTS mapping locations from TTSs in Tiwari et al. show specificity to their designed targets. **A**, HER2-1 (5’-AGGAGAAGGAGGAGGTGGAGGAGGAGGG-3’ in red) connected to all its human genome mapping locations (green) **B**, HER2-205 (5’-GGGGGGCAAGAGGGCGAGGAGGAGCCCC-3’ in red) connected to all its human genome mapping locations (green) **C**, HER2-40118 (5’-GGGGTGGGAGGGACAAAGGGGG-3’ in red) connected to all its human genome mapping locations (green) **D**, HER2-5922 (5’-GGGGTGAGGAGAGTGGGGGAGGAGAAAGGG-3’ in red) connected to all its human genome mapping locations (green)

In the cell line approach, using the BT474 cell line known for ERBB2 amplification (32), the findTTS module identified 18,629 TTSs with 100% precision and accuracy for the cell line’s amplified segments (Figure 5A, B). After further selection of Barcodes indicating the best binding, 2,254 TTSs remained. Clustering on Guanine content and Length reduced this to 53 TTSs with higher Guanine content and length and TTSBBC found a cluster of 207 TTS with better indicators for binding than the HER2-1 cluster (Figure 5D). Another analysis but rather using TTS literature not containing our positive control (15), produced three 100% precise and specific TTSs with optimal binding features with longer lengths than HER2-1, confirming TTSBBC’s effectiveness in diverse data sources (Supplementary Figure 2).

**Figure 5.**
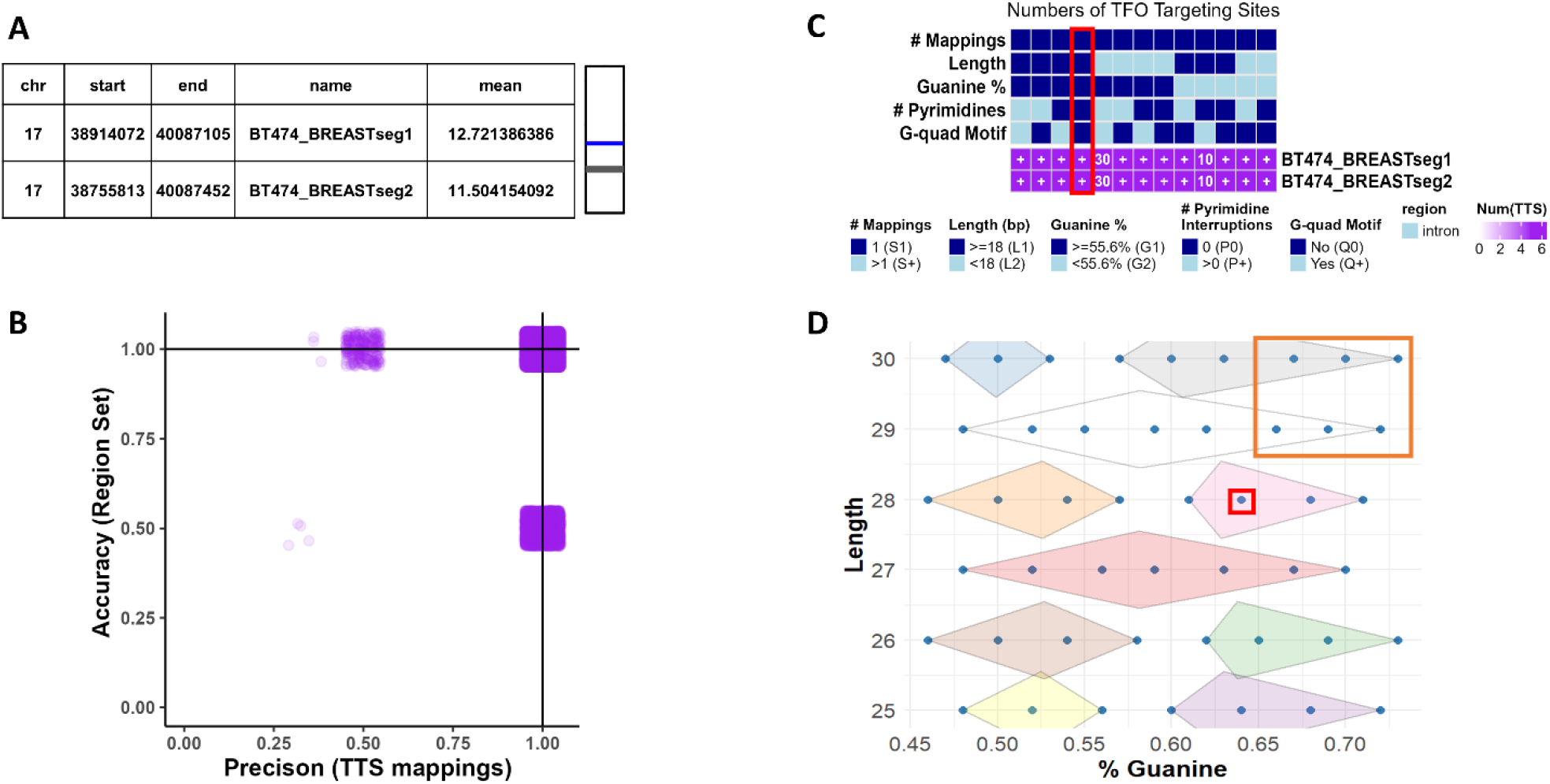
Analyses of TTS in BT474 Amplified Segments reveal putatively better TTS targeting and binding. **A**, Table of amplified segments selected because of high segment means and overlap with ERBB2 to target with TTSs (left) and a visualization of segment location (blue) in relation to the chromosome (right) **B**, Scatterplot (with x and y jitter to show density) of TTS accuracy and precision revealing 18,629 TTSs, of which HER2-1 is one, with 100% accuracy and precision (1,1). **C**, Heatmap of TTS barcodes with query segment targeting. A plus (+) indicates more than 100 TTSs are in that category. HER2-1 is present in the barcodes that indicate the highest binding in the column highlighted in red, along with 2,254 other unique TTSs. **D**, Clustering of TTSs with barcodes that indicate the highest binding (n=2,254) by Guanine content and Length. In the red box is where HER2-1 clustered. In the orange box are TTSs with putatively better TTS targeting and binding (n=53). Grey cluster (n = 207) has better indicators for binding than the cluster that HER2-1 is in (pink).

In the gene-level approach, targeting the ERBB2 gene identified 389 TTS sequences with 100% precision and accuracy (Supplementary Figure 3A). Analysis of their Barcodes (n=60 with ideal Barcodes for binding) and clustering based on Guanine content and Length validated HER2-1 as an effective TTS for targeting ERBB2 with no clear putatively better TTS (Supplementary Figure 3B, C).

This case study found both cell line and gene-based approaches can identify effective TTSs, but targeting amplified segments yielded a larger set of TTSs with better binding features than solely targeting amplified genes. This demonstrates TTSBBC’s utility in guiding researchers to design more effective and efficient experiments targeting any human genomic region, especially those with differential amplification.

*Case Study #2: Discovery of multi-targeting TTSs to the non-homologous end-joining (NHEJ) pathway*. TTSBBC’s design accommodates the fact that a single TTS can have multiple genomic mappings, allowing for the investigation of multiple triplex formations throughout the genome. Triplex structures are known to induce double-stranded breaks (14), necessitating repair via mechanisms like NHEJ. Using the NHEJ pathway as a query in the “findTTS” module, 3,579 TTSs were found to map within NHEJ-related genes. Of these, 6 TTSs showed multi-targeting capabilities with minimal off-target effects (Figure 6A). Analysis of these TTSs’ genic mappings indicated that all 6 mapped to NHEJ1, 2 to XRCC4, 4 to PRKDC, and 6 to ENSG00000280537 (Figure 6B). Barcode screening revealed that none of these TTSs had a G-quadruplex motif, with 2 having high Guanine content and 2 lacking pyrimidine interruptions (Figure 6C). Furthermore, the TTS mappings analysis revealed a high density near NHEJ genes PRKDC, XRCC6, XRCC5, and NHEJ1 (Figure 6D).

**Figure 6.**
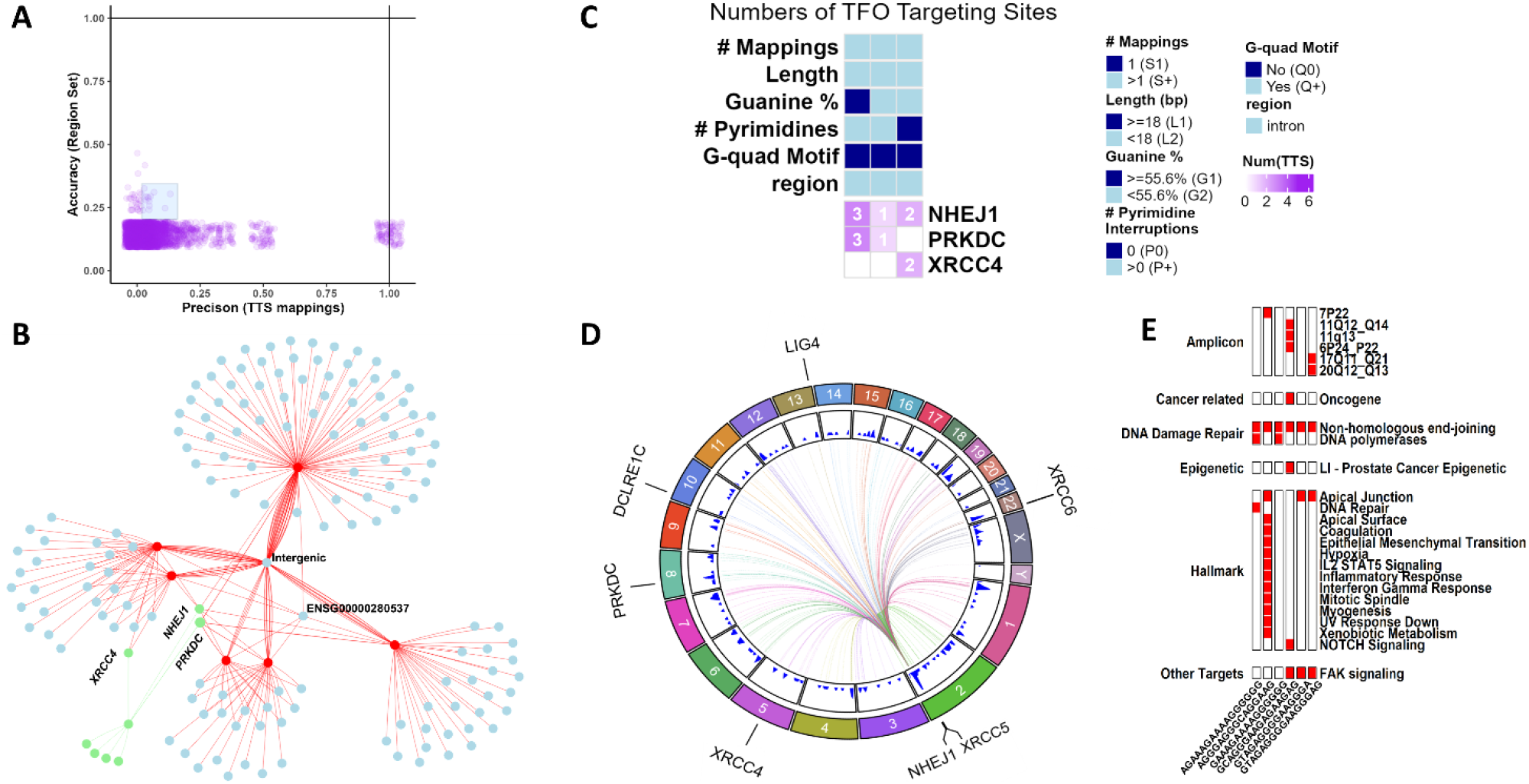
Analysis of non-homologous end-joining (NHEJ) pathway reveals differential multimapping TTS. **A**, Scatterplot (with x and y jitter to show density) of TTS accuracy and precision revealing 6 TTSs, with effective multimapping to query gene set **B**, Network of TTS mapping locations within genic regions. The node colors in red are TTS sequences, in blue are off target genes, and in green are query gene regions. The off-target mappings that are not within any gene region are connected to a blue node labeled intergenic. NHEJ1 contains a mapping from all 6 TTSs, XRCC4 contains 2 of 6, PRKDC contains 4 of 6, and ENSG00000280537 contains 6 of 6. **C**, Heatmap of TTS barcodes with query gene targeting. Query regions, NHEJ1, PRKDC, and XRCC4, contain multiple TTS mapping locations with varying barcodes. **D**, Circos plot of TTS genomic location density. Genomic location density shows concentrations of TTS mappings near PRKDC, XRCC6, XRCC5, and NHEJ1. **E**, Enrichment analysis based on the Hypergeometric distribution (34), using a significance threshold of p≤0.05, of TTS gene signatures as queries and built-in cancer-related gene sets (23-30) as references reveals significant enrichment of TTS gene signatures in red.

Additional analysis revealed off-target effects with complete sequence similarity. However, sequence homology in surrounding regions suggested undiscovered biological mechanisms involving triplex formation possibly involving ENSG00000280537. An enrichment analysis (34) of the TTS gene signature revealed 3 TTSs enriched in FAK signaling, a different set of 3 in Hallmark Apical Junction, 2 in DNA polymerases, and all 6 in NHEJ (Figure 6E). Notably, a single TTS showed enrichment in 12 of the cancer Hallmark gene signatures (26,28). This multi-targeting characteristic of TTSs across the human genome is similar to the DNA-binding domain of transcription factors, which bind to genomic DNA influencing expression and cell differentiation (35,36), and like transcription factors, triplex formation affects transcription (9).

TTSBBC provides a novel approach to study multi-mapping and multi-targeting TTSs, offering unique insights into their biological relationships and mechanisms. This research demonstrates how TTSBBC can help users explore biologic processes and the potential applications of TTS in broader genomic research.

## DISCUSSION

TTSBBC is a comprehensive web platform for analyzing TTSs. It features a user-friendly interface for examining TTS accuracy, precision, Barcodes, genomic mappings, and sequence-level characteristics. The platform highlights the multi-targeting potential of TTSs, in addition to their traditional single targeting applications, offering novel insights into the biological and therapeutic implications of their multi-mapping nature.

TTSBBC, specializing in human genomic analysis of TTSs, differs from other methods that focus on TFO design and assessment (37,38). Unlike Triplex-Inspector, which assesses TFO-TTS conjugates and their off-target effects (39,40) while ignoring non-specific TTSs, TTSBBC focuses assessments of specific and non-specific TTSs with analyses capabilities. TTS Mapping, finds TTSs for specific regions but restricts focus to highly-specific TTSs and co-occurring motifs (41). Notably, at the time of writing, these tools are not accessible via their published links. TFO-Search and TTSMI are TTS sequence repositories without analysis features (15,16), and are included in TTSBBC as data sources. TTSBBC advances beyond these tools by offering analytical capabilities with informed and smart sequence screening for experimental design.

Future planned enhancements for TTSBBC include the ability to analyze sets of TTSs, their genomic mappings, and features to aid in precision medicine and personalized anti-gene therapies. Another potential improvement is incorporating all possible permutations of TTS sequences, not just those from published sources, for cartography of the landscape of TTSs and their multi-targeting potential. The initial curation of published and experimentally validated TTSs remains ongoing and will be updated, potentially forming a third database in the future. A forthcoming update includes TTS mapping to the newest Telomere-to-Telomere (T2T) (42) coordinates and annotations.

The web server offers various input options to enhance user experience and is accessible to a wide audience. Its intuitive visualizations aid non-computational biologists in exploring TTS sequence features and genomic mappings for targeted genomic instability in cancer. TTSBBC introduces novel adaptations of metrics and concepts for TTSs - precision, accuracy, Barcoding, and multi-targeting assessment – integrating these into a comprehensive and accessible analysis platform.

## DATA AVAILABILITY

The TTSBBC app (https://kowalski-labapps.dellmed.utexas.edu/TTSBC/) is a freely available web server that does not require a login.

## SUPPLEMENTARY DATA

Supplementary Data are available at NAR online.

## AUTHOR CONTRIBUTIONS

M.Y. performed the analyses, drafted the figures and initial manuscript. M.Y. and Q.X. designed the platform and implemented the workflows. J.K. conceived of the idea, directed the analyses, and edited the manuscript.

## ACKNOWLEDGEMENT

The website of TTSBBC is hosted on an AWS server provided by the Kowalski Lab and is supported by UT Austin IT solutions and Dell Medical School at UT-Austin.

