## Supplemental Data 1 for "TTSBBC: Triplex Target Site Biomarkers and Barcodes in Cancer": README.txt

################
###  README  ###
################

This directory contains 2 files:
- tfo_search_bc_sig.txt
- ttsmi_bc_sig.txt

These are the Barcode signatures for each respective datasource. The format for each file has each Barcode's signature on a line.  For a given line, the Barcode is the first piece of text, followed by a tab("\t"), followed by a comma-separated list of gene symbols.  If a gene symbol is not available an Ensembl gene code is used.

For example:
G1_L2_S+_P+_Q0	TP73,AJAP1,RPL22

or more generally for N genes:
<Barcode>\t<Gene1>,<Gene2>,<Gene3>,...,<GeneN>

These genes are extractable with common text manipulation software such as regular expressions or in Excel using tab as a delimiter
