## Supplemental Figures for "TTSBBC: Triplex Target Site Biomarkers and Barcodes in Cancer"

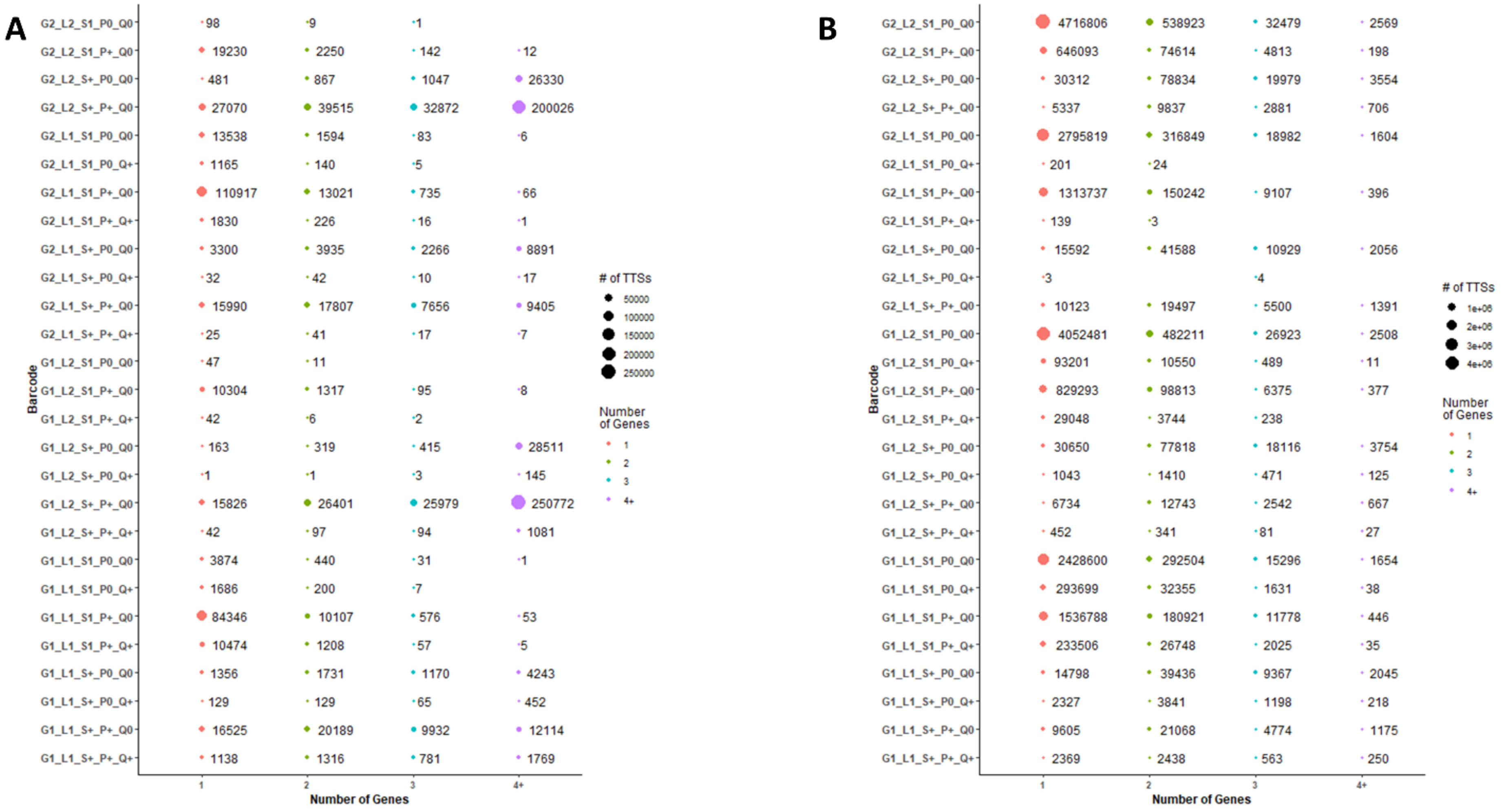


Supplemental Figure 1. **TTS gene specificity relative to barcodes illustrates patterns of genic specificity.** The locations of TTSs with the provided barcode were joined with gene regions to find the gene signature for the TTS. The sizes of the TTS gene signature were then plotted against the barcode.

**A**, Barcodes and TTSs from the TFO-Search data source

**B**, Barcodes and TTSs from the TTSMI data source

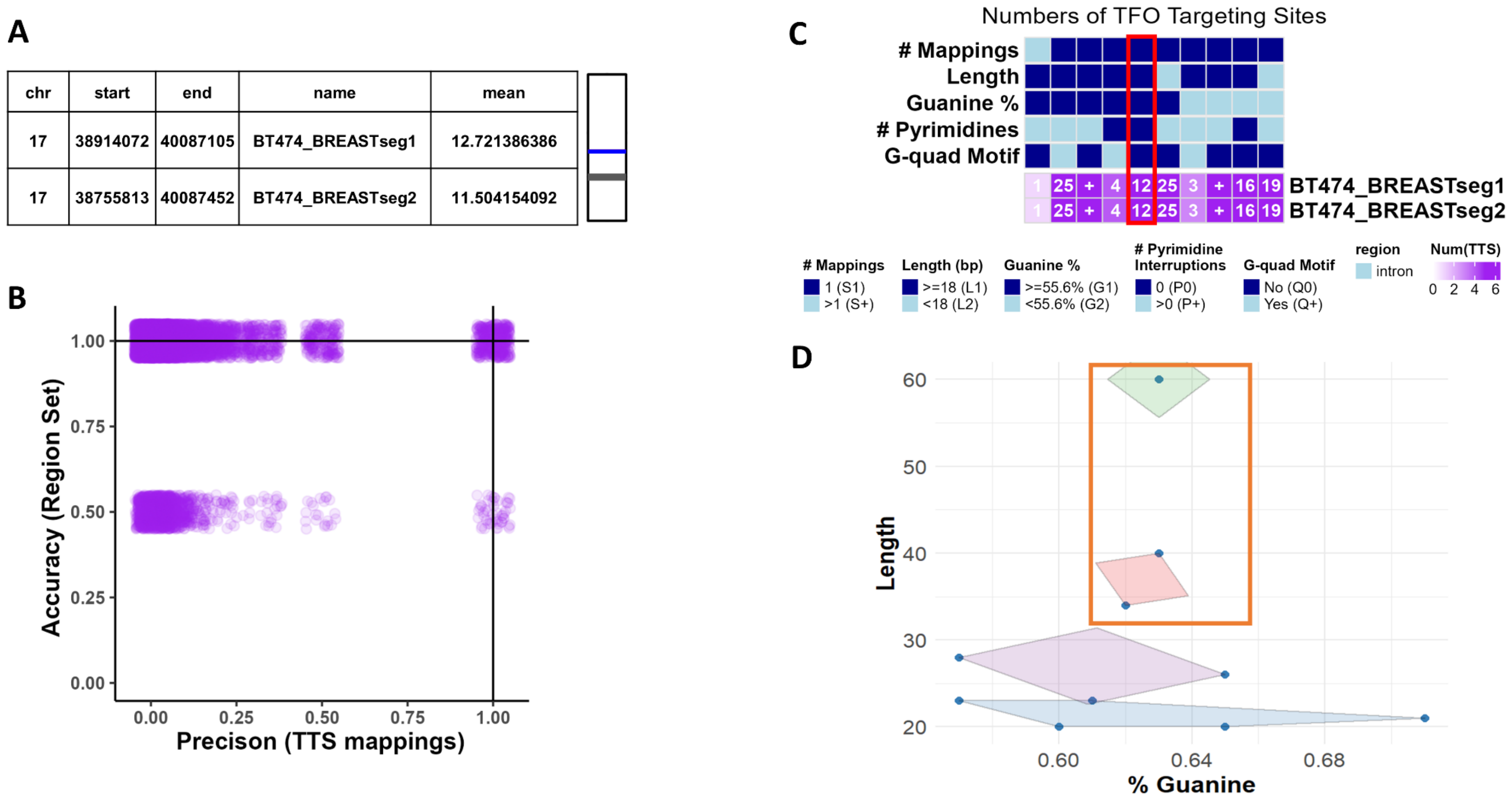


| chr | start | end | name | mean |
|---|---|---|---|---|
| 17 | 38914072 | 40087105 | BT474_BREASTseg1 | 12.721386386 |
| 17 | 38755813 | 40087452 | BT474_BREASTseg2 | 11.504154092 |

Supplementary Figure 2. **Analyses of TTS in BT474 Amplified Segments reveal putatively better TTS targeting and binding**.

**A**, Table of amplified segments selected because of high segment means and overlap with ERBB2 to target with TTSs (left) and a visualization of segment location (blue) in relation to the chromosome (right)

**B**, Scatterplot (with x and y jitter to show density) of TTS accuracy and precision revealing 375 TTSs, with 100% accuracy and precision (1,1).

**C**, Heatmap of TTS barcodes with query segment targeting. A plus (+) indicates more than 100 TTSs are in that category. The barcodes that indicate the highest binding in the column highlighted in red (n=12).

**D**, Clustering of TTSs with barcodes that indicate the highest binding (n=12) by Guanine content and Length. In the orange box are TTSs with putatively better TTS targeting and binding (n=3).

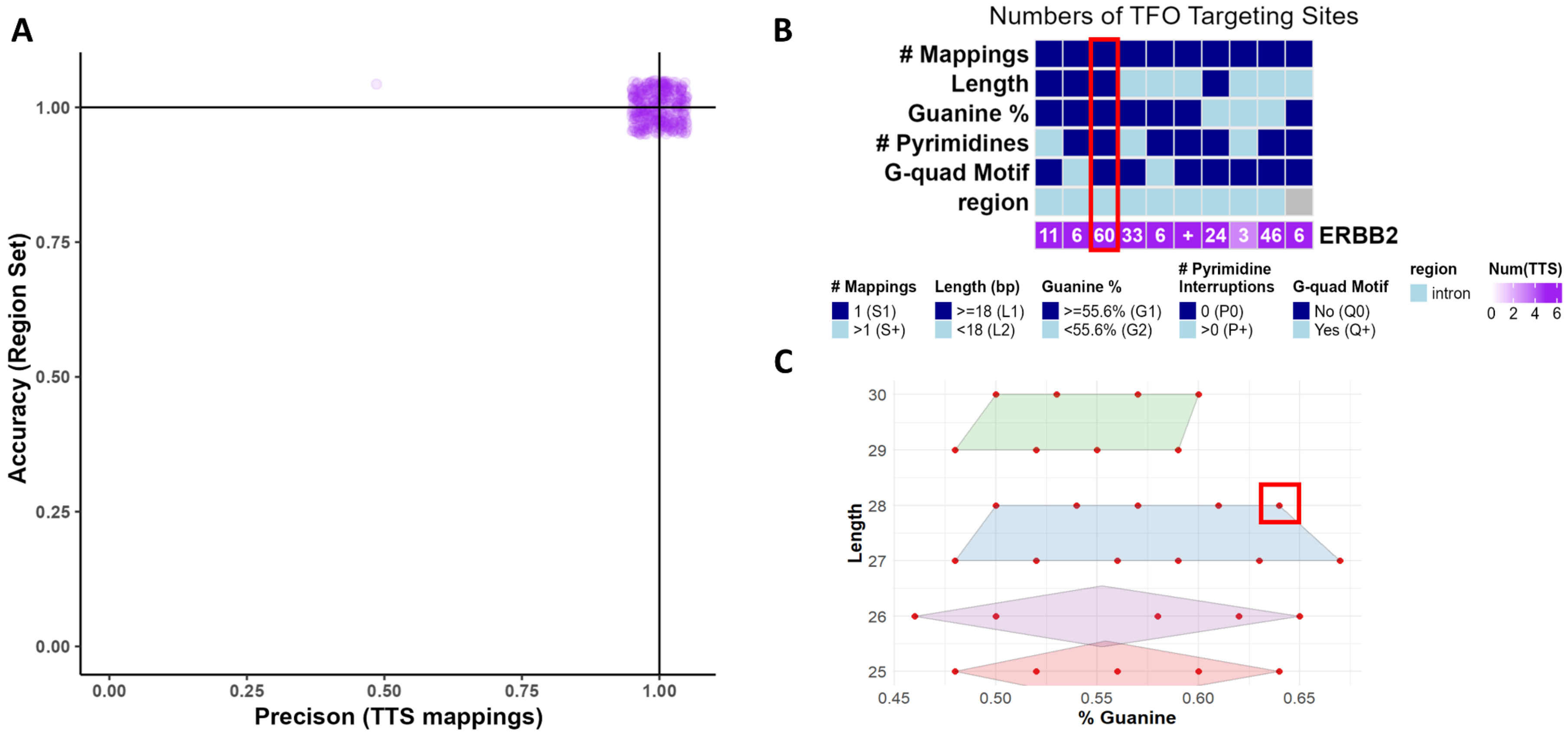


Supplementary Figure 3. **Analyses of TTS in ERBB2 reveal TTS selection quality**.

**A**, Scatterplot (with x and y jitter to show density) of TTS accuracy and precision revealing 389 TTSs, of which HER2-1 is one, with 100% accuracy and precision (1,1).

**B**, Heatmap of TTS barcodes with query segment targeting. A plus (+) indicates more than 100 TTSs are in that category. HER2-1 is present in the barcodes that indicate the highest binding in the column highlighted in red, along with 60 other unique TTSs.

**C**, Clustering of TTSs with barcodes that indicate the highest binding (n=60) by Guanine content and Length. In the red box is where HER2-1 clustered.
